# Investigating the Osteoinductive Potential of a Decellularized Xenograft Bone Substitute

**DOI:** 10.1101/419101

**Authors:** Daniel N. Bracey, Alexander H. Jinnah, Jeffrey S. Willey, Thorsten M. Seyler, Ian D. Hutchinson, Patrick W. Whitlock, Thomas L. Smith, Kerry A. Danelson, Cynthia L. Emory, Bethany A. Kerr

## Abstract

Bone grafting is the second most common tissue transplantation procedure worldwide. The gold standard for bone grafting is the autograft; however, due to morbidity and limited supply, new alternatives, including allograft and tissue-engineered bone substitutes, are needed to satisfy long-term demand. One of the most desired properties of tissue-engineered bone substitutes is osteoinductivity, defined as the ability to stimulate primitive cells to differentiate into a bone forming lineage. In the current study, we treated porcine bone with a decellularization protocol to produce a bone scaffold. We examined whether the scaffold possessed osteoinductive potential and could be used to create a tissue-engineered bone microenvironment. To test if the bone scaffold was a viable host, pre-osteoblasts were seeded, incubated *in vitro*, and analyzed for markers of osteogenic differentiation. To assess these properties *in vivo*, scaffolds with and without pre-osteoblasts pre-seeded were subcutaneously implanted in mice for four weeks. The scaffolds underwent micro-computed tomography (microCT) scanning before implantation. After retrieval, the scaffolds were analyzed for osteogenic differentiation or re-scanned by microCT to assess new bone formation with the subsequent histological assessment. The osteoinductive potential was observed *in vitro* with similar osteogenic markers being expressed as observed in demineralized bone matrix and significantly greater expression of these markers than controls. By microCT, paired t-tests demonstrated significantly increased bone volume:total volume (BV/TV) and trabecular thickness (Tb.Th) after explantation in all groups. Pentachrome staining demonstrated osteogenesis within the scaffold, and angiogenesis in the scaffold was confirmed by CD31 staining for blood vessels. These results demonstrate that porcine bone maintains its osteoinductive properties after the application of a novel decellularization and oxidation protocol. Future work must be performed to definitively prove osteogenesis of human mesenchymal stem cells, biocompatibility in large animal models, and osteoinduction/osseointegration in a relevant clinical model *in vivo*. The ability to create a functional bone microenvironment using decellularized xenografts will impact regenerative medicine, orthopaedic reconstruction, and could be used in the research of multiple diseases.

## 2. Introduction

Regeneration and healing of critical bone defects resulting from high energy trauma, infection, necrosis, or tumor resection remain a major clinical challenge for orthopaedic surgeons [Calori et al., 2011; Kolambkar et al., 2011; Fassbender et al., 2014; Oryan et al., 2017]. The local biology in these defects is disrupted, rendering the self-regenerative healing cascade insufficient, and thus, conventional reparative techniques lead to nonunions, malunions, and osteomyelitis [Oryan et al., 2014; Oryan et al., 2017]. The gold standard treatment for a traumatic bone defect is the use of autologous bone graft; however, due to the associated morbidity and lack of adequate bone stock/donor sites, alternative grafts are commonly used. Alternative bone grafts include allografts and tissue-engineered bone substitutes [Calori et al., 2011; Roddy et al., 2018]. Allograft use risks disease transmission and has limited availability from young, healthy donors [De Long et al., 2007; Campana et al., 2014]. Consequently, there is increased interest in tissue-engineered bone substitutes [Wanschitz et al., 2007; Pina et al., 2017; Shahi et al., 2018; Iaquinta et al., 2019].

The ideal tissue-engineered bone substitute will be osteoconductive, osteoinductive, and osteogenic [Calori et al., 2011; Oryan et al., 2017]. Osteoconductivity is the ability for the bone graft to allow osseous growth on the surface or within its pores [Khan et al., 2005]. Osteogenic grafts retain living bone cells [Khan et al., 2005]. Osteoinductivity is the ability to stimulate progenitor cells to differentiate into a bone forming cell lineage [Albrektsson and Johansson, 2001; Khan et al., 2005]. Tissue-engineered bone replacements can be manufactured with customized structures for osteoconductivity and pre-seeded with osteogenic cells to establish osteogenicity prior to implantation [Zimmermann and Moghaddam, 2011]. Osteoinductivity, however, requires the construct to induce cell differentiation and therefore is more difficult to recreate [Albrektsson and Johansson, 2001; Fielding and Bose, 2013; Hsu et al., 2013; Oryan et al., 2017]. Xenograft derived tissue-engineered constructs are one potential way to utilize the natural osteoinductive properties of native bone. One ideal species for xenotransplantation is swine, due to physiologic compatibility with humans [Pierson et al., 2009; Wancket, 2015]. However; the presence of the alpha-gal epitope in porcine tissue can induce a severe inflammatory response in human hosts [Cooper et al., 2015; Vadori and Cozzi, 2015]. Our laboratory developed a decellularization protocol that sterilizes porcine soft tissues and removes the porcine DNA, including the alpha-gal epitope [Whitlock et al., 2007; Whitlock et al., 2012; Seyler et al., 2017]. We have applied this process to porcine cancellous bone and demonstrated that the construct was successfully decellularized and maintained native structural properties, therefore preserving the construct’s osteoconductivity [Bracey et al., 2018]. This research project aimed to determine whether the osteoinductive potential of the porcine-derived bone scaffold would be maintained following application of a novel decellularization and oxidation technique in *in vitro* and in vivo *models*.

## 3. Materials and Methods

### Bone Scaffold Decellularization

To generate the bone scaffolds, porcine tissues were processed by decellularization, and oxidation using methods previously described [Whitlock et al., 2012; Seyler et al., 2017; Bracey et al., 2018]. Briefly, the cancellous bone was harvested from the distal metaphysis of porcine femurs and subjected to chemical decellularization and oxidation using combinations of deionized water, trypsin, antimicrobials, peracetic acid (PAA), and triton-X. Scaffolds were lyophilized and frozen at −80°C until further use. Scaffolds were cut to 1 cm diameter, and the thickness ranged between 0.2 to 0.5 cm. The bone scaffolds were not decalcified at any point during this process.

### Cell Culture

C2C12 and MC3T3-E1 cell lines were chosen for indirect quantification of the bone scaffold’s osteoinductive potential. C2C12 (CH3 genetic background, ATCC^®^ CRL1772™, Rockville, MD) is a mouse myoblast cell line that differentiates into osteoblasts in the presence of BMP-2 and is commonly used in osteoinduction studies [Katagiri et al., 1994; Han et al., 2003; Yang et al., 2011; Shi et al., 2012; Ansari et al., 2013]. C2C12 cells were grown in DMEM media supplemented with 10% FBS. MC3T3-E1 cells (ATCC^®^ CRL1772™, Rockville, MD) were chosen as a second cell line to confirm the bone scaffold’s osteoinductive potential. These cells are an osteoblast precursor derived from C57BL/6 mice and have previously been used to study osteoinductive potential as well [Shuang et al., 2016; Araujo-Gomes et al., 2018]. MC3T3-E1 cells were grown in αMEM media supplemented with 10% FBS and sodium pyruvate.

### Osteogenic Differentiation of C2C12 Cells

Bone scaffolds or commercial grade cancellous demineralized bone matrix (DBM, Musculoskeletal Transplant Foundation, Edison NJ) sheets (n=77 per group) were seeded with 1×10^6^ C2C12 cells suspended in 100 μl cell culture media. DBM is commonly used in clinical applications because of its reported osteoconductive and osteoinductive potential. Constructs were moved into large Petri dishes, covered with DMEM + 10% FBS media, and returned to the incubator. As a negative control, cells were also seeded onto gelfoam sponges (Cardinal Health) which are assumed to have no or very limited biologic activity.

Constructs were incubated for 24 hours to allow cells to attach to the matrix. Sub-samples from each group (n=11) were taken for analysis while the remaining were separated for continued incubation in 2 different medias: 1) “Osteogenic Media (OM)” [Shui et al., 2013; Sondag et al., 2013; Yu et al., 2013; Hupkes et al., 2014] consisting of DMEM with 10 mM β-Glycerophosphate and 50 μg/mL ascorbic acid or 2) “BMP-2 Enriched Media” [Han et al., 2003; Feichtinger et al., 2011; Yang et al., 2011; Ansari et al., 2013] consisting of the osteogenic media supplemented with 100 ng/mL BMP-2 (recombinant human BMP-2, 355-BM-050, R&D Systems). Media was replaced every 3 days. The osteogenic media provided an environment supportive of osteogenic differentiation while the BMP-2 enriched media served as a positive control to drive cells towards osteoblastic lineage. Constructs (n=11) were harvested from each group at days 1, 3, 7, and 15 for analysis of cell proliferation and osteogenic differentiation.

### Cell Viability and Proliferation on Scaffolds

At each time point, constructs (n=2) from each group (n=6) were removed from their respective dishes and individually rinsed with warm, sterile PBS. Constructs were then transferred to chamber slides and incubated with the Live/Dead^®^ Viability/Cytotoxicity Kit (Molecular Probes, Eugene, OR) according to manufacturer’s instructions. Specimens were immediately imaged on a fluorescent confocal microscope (Zeiss Axiovert 100 M) to render cross-sectional 2D images as well as projected 3D images. Live cells are labeled with the green calcein AM fluorophore, and dead cells are labeled with the red ethidium homodimer-1 fluorophore.

DNA content was quantified from separate constructs (n=3) in each group to estimate cell number and proliferation. Samples were flash frozen in liquid nitrogen, homogenized with a sterilized tissue press, and lysed in 1 mL mammalian protein extraction reagent (M-PER^®^, ThermoScientific, Waltham, MA). Samples were centrifuged 15 minutes at 13.2 kRPM and supernatants were collected for analysis. DNA content was measured with Quant-iT™ PicoGreen^®^ dsDNA Assay Kit (Thermo Scientific) according to the manufacturer’s instructions.

### Scanning Electron Microscopy and Scaffold Histology

Cell attachment, morphology, and surface distribution were characterized by electron microscopy. Specimens (n=2) were removed from dishes in each group at every time point, gently washed with warmed sterile PBS and fixed in 2.5% glutaraldehyde for 3 hours and imaged on a Hitachi S-2600 scanning electron microscope (SEM).

Specimens (n=2) from each group were removed from dishes, fixed in 10% formalin for 48 hours, decalcified with Immunocal^®^ (Decal Chemical Cort, Tallman, NY) for 3-5 days, processed and embedded in paraffin. Sections were mounted and stained with hematoxylin and eosin (H&E), Masson’s trichrome or 4’,6-diamidino-2-phenylindole (DAPI) mounting media (ProLong^®^ Gold Antifade Mountant, Thermo Scientific). Representative light micrographs were captured with the Olympus VS-110 Virtual Imaging System and fluorescent micrographs with a Zeiss Axioplan2 system.

### Alkaline Phosphatase Enzyme Assay and Immunohistochemistry

The alkaline phosphatase (ALP) enzyme activity was measured from individual constructs (n=3) using methods previously described [Liu et al., 2008; Stiehler et al., 2010; Thibault et al., 2010; Arca et al., 2011; Marcos-Campos et al., 2012; Shi et al., 2013]. Each reaction was read in triplicate by loading 150 μl from each tube into 96 well plates and measuring absorbance at 405 nm on a microplate reader (Spectra Max 340 PC). Cell-specific ALP activity was determined by normalizing enzyme activity to the respective sample’s DNA content determined by Pico Green assay [Arca et al., 2011; Hashimoto et al., 2011; Kouroupis et al., 2013].

Immunohistochemistry (IHC) was performed with an anti-Placental ALP (Abcam ab16695, Cambridge, United Kingdom) primary antibody that reacts with cell membrane-bound enzyme. Secondary biotin-conjugated anti-rabbit antibody (BioGenex, Freemont, CA) was linked to HRP (horseradish peroxidase) and developed with DAB (diaminobenzidine) substrate (Vector, Burlingame, CA). Slides were counterstained with Mayer’s Hematoxylin.

### Differentiation of MC3T3-E1 cells

MC3T3-E1 cells (1×10^6^) suspended in 250 μL of cell culture media were seeded and incubated on scaffolds (n=18) for 1 hour before being submerged in 750 μL of α-MEM and incubated at 37° C for 7 days. Control monolayers (n=9) of 1 x 10^6^ cells plated on 10 μg/mL collagen type I (Sigma) were also incubated at 37° C for 7 days in α-MEM. Cell culture media was changed every 3 days.

### qPCR Analysis

RNA was isolated from the scaffolds by grinding in liquid nitrogen followed by lysis in Qiazol Reagent (RNeasy Microarray Tissue Mini Kit, QIAGEN, Hilden, Germany), and then RNA purified following the manufacturer instructions for. RNA from monolayers was also isolated following manufacturer instructions using the RNeasy Mini Kit. cDNA was produced through reverse transcription using High Capacity cDNA Reverse Transcription Kit (Applied Biosystems) and then analyzed for gene expression of different osteoblast markers using quantitative PCR. Target gene expression was normalized to the housekeeping gene 18S ribosomal RNA (rRNA). Primer sequences are available in Table 1. Relative gene expression was quantified by using the 2^-ΔΔct^ methodology originally described by Livak and Schmittgen [Livak and Schmittgen, 2001].

**Table 1.**
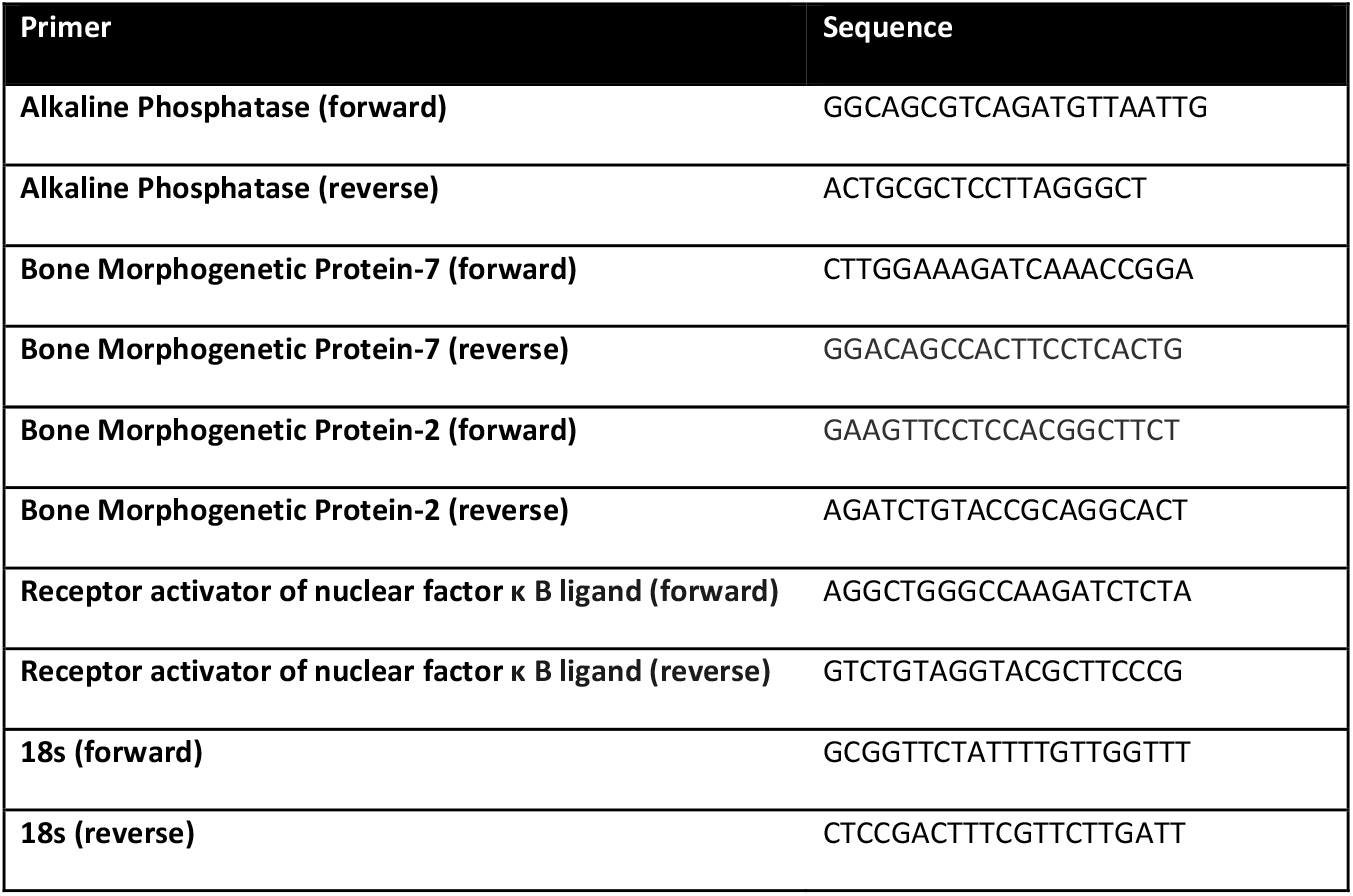
qPCR primer sequences

### Subcutaneous Implantation of Scaffolds

To demonstrate cell viability and osteoinductivity *in vivo*, the scaffolds were either preseeded (n=25) with MC3T3-E1 cells (1 x 10^6^ suspended in 50 μL α-MEM) or not pre-seeded with any cells (n=15). These scaffolds were then subcutaneously implanted in 7-week-old C57BL/6 mice (Jackson Labs, Bar Harbor ME) for 4 weeks under a Wake Forest School of Medicine IACUC Protocol #A16-197 (one scaffold per mouse).

### Micro-computed Tomography (microCT)

A subset of scaffolds (n=12 pre-seeded, n=7 unseeded) underwent micro-computed-tomography (microCT) scanning (TriFoil Imaging Triumph PET/CT, voxel size 1mm) before cell seeding and implantation. Upon explantation, the constructs were placed in Buffer RLT (QIAGEN) and vortexed to extract RNA or placed in 10% formalin to undergo microCT scanning to assess new bone formation. For scaffolds that underwent microCT scanning, new bone formation was assessed through the change in bone volume/total volume ratio (BV/TV) and trabecular thickness (Tb.Th) using MicroView 3D Image Viewer and Analysis Tool (Parallax Innovations, Ilderton, ON, Canada).

### Implanted Scaffold Histology and Immunohistochemistry

Samples that underwent microCT scanning were fixed in 10% formalin overnight on a shaker at 4°C and subsequently decalcified in 14% neutral, saturated EDTA for 7-14 days. Samples were processed and embedded in paraffin. Sections were stained with Russel-Movat-Pentachrome (American MasterTech Scientific Inc; St. Lodi, CA). IHC was performed using primary antibodies against osteopontin (OPN) (Abcam ab8448), ALP (R&D Systems FAB1448A) and CD31 (Abcam ab28364) to identify active bone remodeling, new bone formation, and angiogenesis. Slides were scanned using a Hamamatsu NanoZoomer by the Virtual Microscopy Core.

### Statistical Analysis

Multiple group comparisons were performed using one-way ANOVA, t-tests were performed on independent means when comparing two groups, and paired t-tests were used when comparing paired groups. Statistical significance was determined when α-error<0.05. Graphs were prepared in GraphPad Prism 7.

## 4. Results

### Scaffolds Support C2C12 Pre-Osteoblast Attachment, Survival, and Proliferation

C2C12 pre-osteoblasts proliferated on scaffolds and deposited extracellular matrix (ECM) components. Three-dimensional images of Live/Dead staining demonstrated circumferential cell attachment evenly around the pores. In all constructs, the live (green) signal increased during incubation with the strongest signal noted at day 15 (Figure 1). Dead (red) signal was strongest on day 1, likely due to early contact inhibition after seeding constructs at a high cell density. Additionally, all constructs displayed autofluorescence in the red channel as shown in the Blank images (Figure 1). To assess proliferation on the constructs, DNA content was measured on the three constructs over 15 days in the presence or absence of 100 ng/mL BMP-2 to induce osteoblast differentiation. DNA content on DBM was greater than scaffolds at every time point, indicating higher cell density (Figure 2). BMP-2 treatment had no consistent effect on cell proliferation. Cells seeded on the scaffold proliferated, as demonstrated by increased DNA at each time point with a 6.5-fold increase between days 1 and 15. The differences were significant between day 1 and 15 (p<0.01) and day 3 and 15 (p=0.01). Similarly, DNA content on DBM increased at each time point as well (Figure 2), while the gelfoam constructs demonstrated increased DNA content at day 7 with a decrease at day 15. To confirm cell numbers and adhesion to the constructs, C2C12 seeded DBM and decellularized bone scaffolds were examined. DAPI and H&E staining (Figure 3) confirmed higher cell density on DBM relative to scaffolds indicating that the increased cell proliferation is occurring in cells attached to the constructs. Finally, scanning electron microscopy confirmed C2C12 adhesion to and spreading on the constructs. Dense cell distribution was noted on DBM samples relative to scaffolds at day 7 and 15 (Figure 4). ECM was deposited uniformly at later time points, and BMP-2 did not change cell morphology, density, or distribution on matrices. Thus, the decellularized bone scaffolds were equally capable of supporting pre-osteoblast adhesion and survival underscoring the osteogenic nature of the scaffolds.

**Figure 1.**
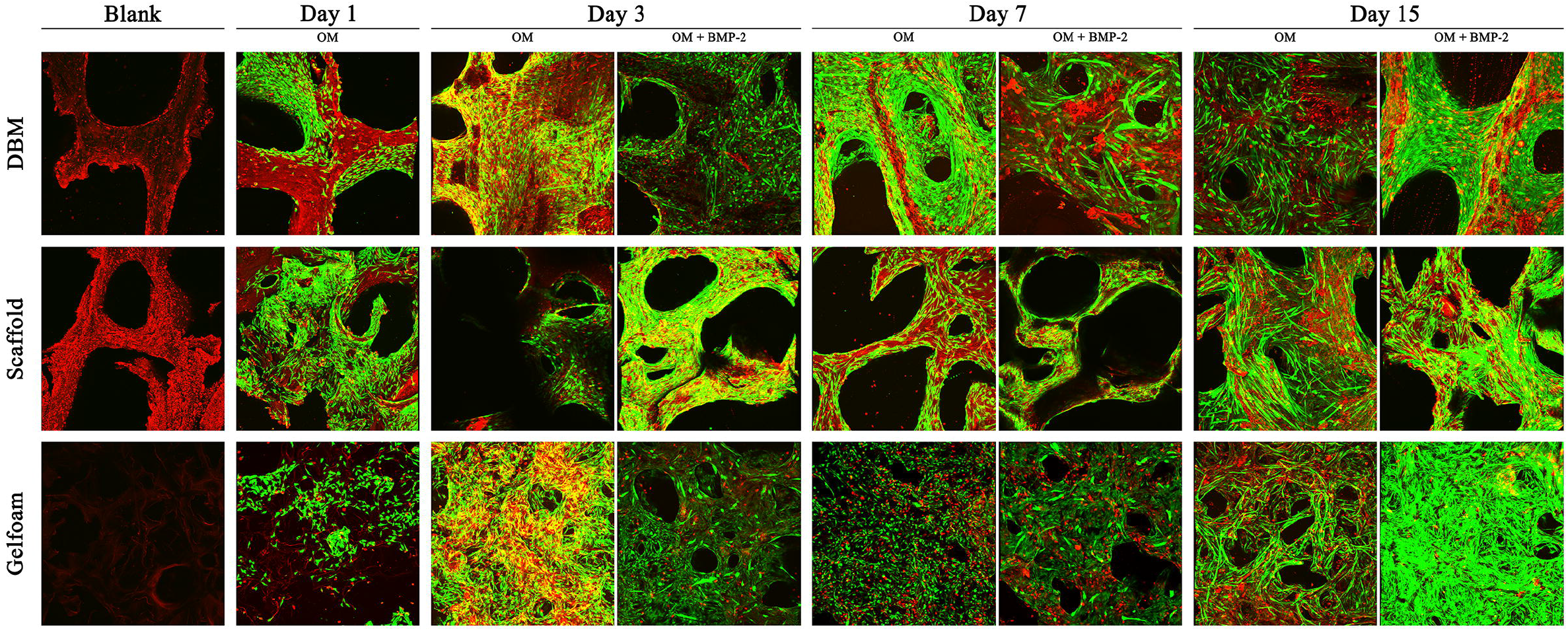
C2C12 cells are viable on the decellularized bone scaffold. Demineralized Bone Matrix (DBM), scaffold, and gelfoam matrices were seeded with 1 million C2C12 pre-osteoblast cells, incubated for 1, 3, 7, or 15 days. On day 1, matrices were switched into osteogenic media (OM) enriched with 100 ng/mL BMP-2. Representative micrographs show green fluorophore (calcein AM) staining of live cells and red fluorophore (ethidium) staining of dead cells. Constructs had notable autofluorescence with ethidium staining as shown in the “blank” images. Cross-sectional images were captured at 10X magnification and overlayed to create the shown 3D projections. Cell density increased with time on all 3 matrices, consistent with DNA quantification results.

**Figure 2.**
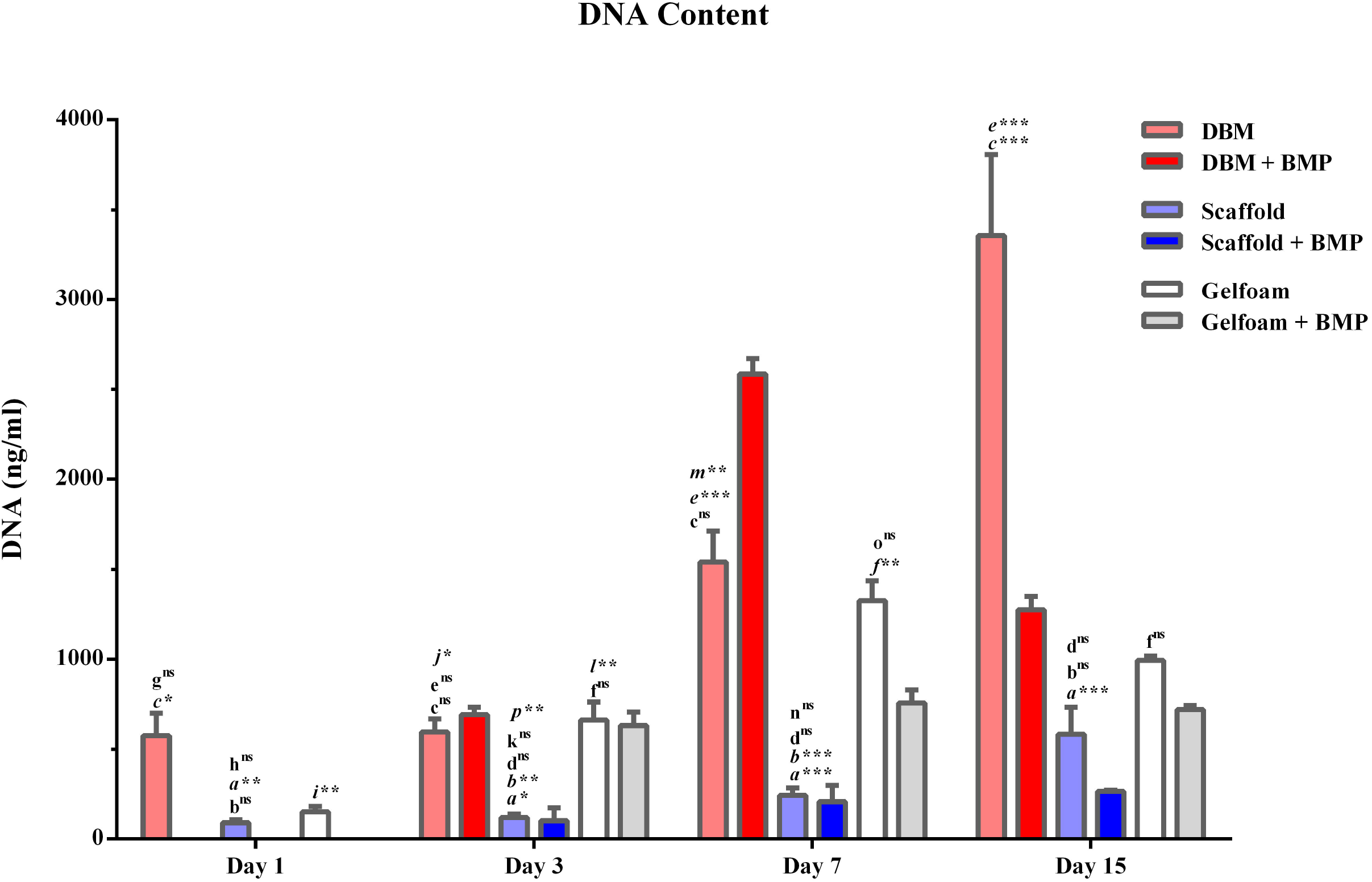
C2C12 cells proliferate slowly on the decellularized bone constructs. DBM (red bars), bone scaffold (blue bars), or gelfoam (GF; white bars) constructs were harvested at days 1, 3, 7, and 15, and treated with control media, osteogenic media (OM), or OM with 100 ng/mL BMP-2. DNA content was measured with the PicoGreen^®^ assay and represented as mean DNA content ± SEM. * represents p<0.05, ** represents <0.01, and *** represents p<0.001. **(a)** scaffold vs DBM; **(b)** scaffold vs GF; **(c)** DBM vs GF **(d)** scaffold OM vs BMP; **(e)** DBM OM vs BMP; **(f)** GF OM vs BMP; **(g)** DBM day 1 vs 3; **(h)** scaffold day 1 vs 3; **(i)** GF day 1 vs 3; **(j)** DBM day 3 vs 7; **(k)** scaffold day 3 vs 7; **(l)** GF day 3 vs 7 **(m)** DBM day 7 vs 15; **(n)** scaffold day 7 vs 15; **(o)** GF day 7 vs 15 **(p)** scaffold day 3 vs 15; **(q)** DBM day 3 vs 15; **(r**) GF day 3 vs 15

**Figure 3.**
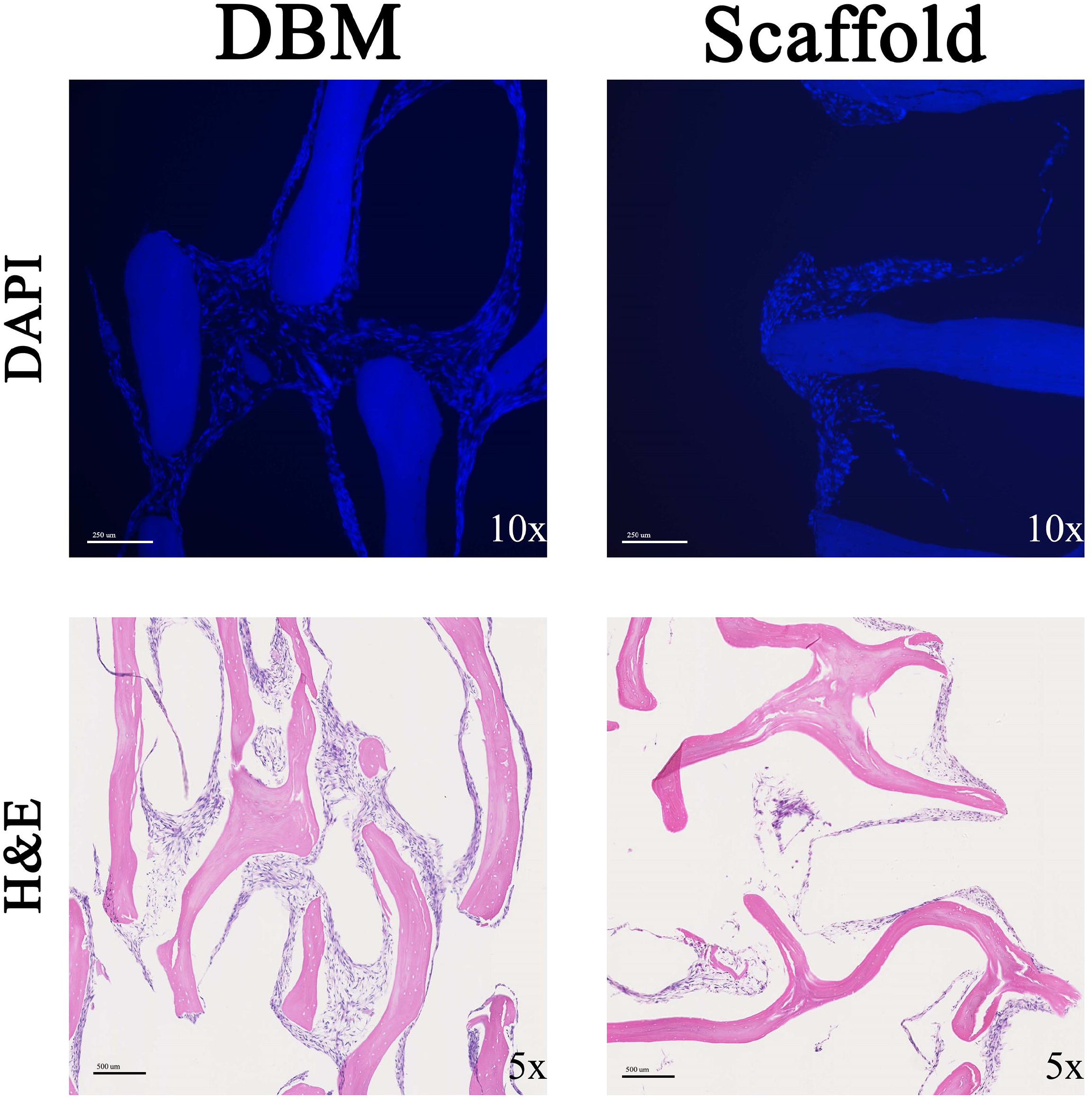
C2C12 cell density is increased on DBM compared with scaffolds. C2C12 cells were seeded on DBM (left panels) or decellularized bone scaffold (right panels). Seeded scaffolds were sectioned and stained for DAPI (top panels) or H&E (bottom panels). Representative images are shown taken at 10X (top panels) or 5X (bottom panels).

**Figure 4.**
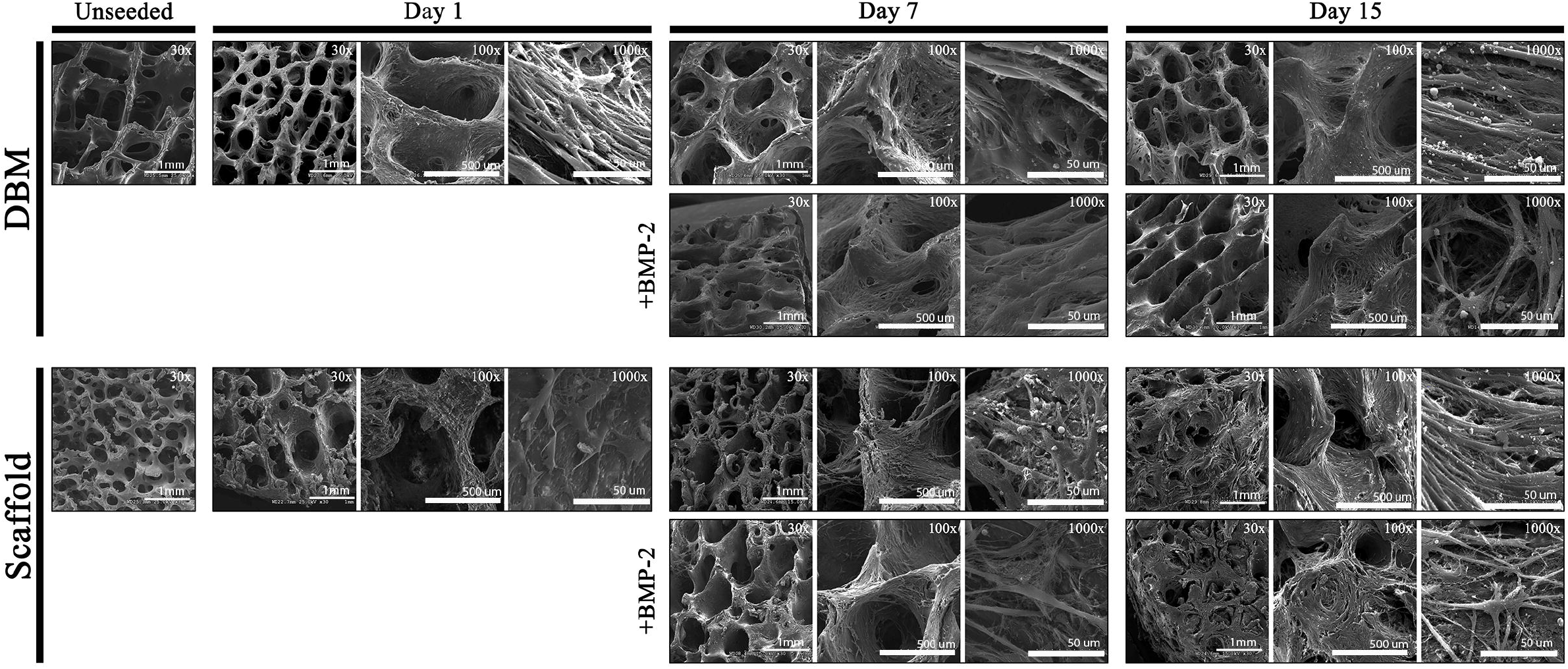
C2C12 attachment and spreading is equivalent on bone constructs and DBM. C2C12 seeded DBM and bone scaffolds were analyzed by scanning electron microscopy at days 1, 7, or 15 and compared to unseeded scaffolds. Comparison with 100 ng/mL BMP-2 treated scaffolds are shown. Representative images at 30X (scale bar represents 1 mm), 100X (scale bar represents 500 μm), and 1000X (scale bar represents 50 μm) are shown.

### Decellularized Bone Scaffolds Enhance Pre-Osteoblast Differentiation

To examine the osteoinductive capacity of the decellularized bone scaffolds, C2C12 preosteoblast differentiation on the constructs was compared. C2C12 cells were seeded onto either DBM, bone scaffolds, or Gelfoam and grown in the presence or absence of BMP-2 for 15 days.

Molecular assays demonstrated that cells seeded on decellularized bone scaffolds had greater ALP enzyme activity at day 7 (~40-fold) and day 15 (~25-fold) (p<0.0001) compared to cells seeded on DBM or gelfoam constructs (Figure 5). BMP-2 increased ALP activity on all constructs, which was only significant for scaffolds (days 3 (p=0.005), 7 (p=0.02), 15 (p<0.0001)) suggesting an additive effect on this matrix. ALP IHC staining increased at day 15 in C2C12 cells seeded on bone scaffolds and supported the above cell-specific enzyme activity findings (Figure 6). ALP staining demonstrates darker staining on the scaffolds compared with DBM in the presence or absence of BMP-2. Thus, the bone scaffolds support osteoblast differentiation of C2C12 cells.

**Figure 5.**
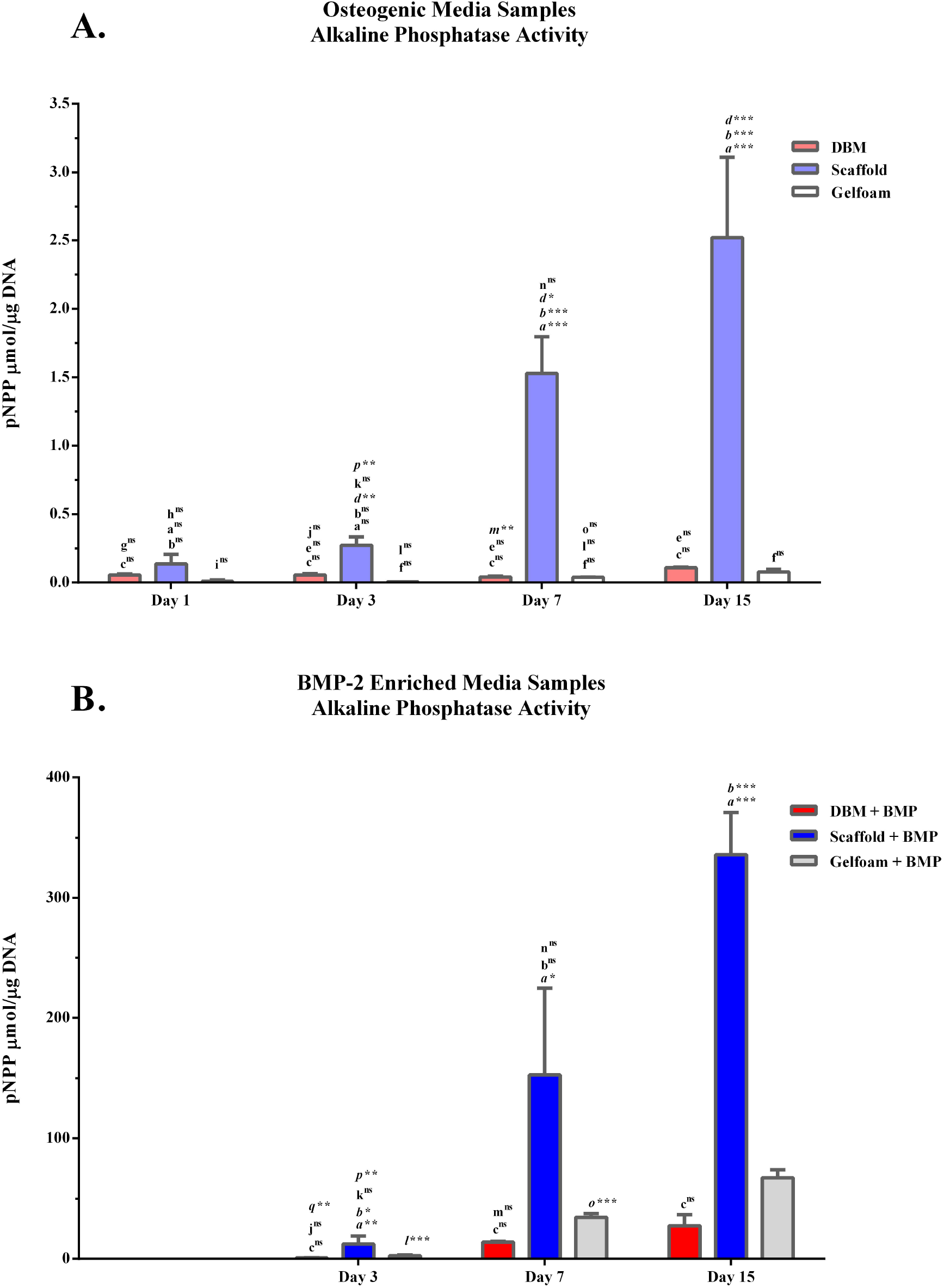
Scaffolds induce ALP activity in C2C12 cells. C2C12 seeded constructs: DBM (red bars), bone scaffolds (blue bars), or Gelfoam (white bars), were incubated in osteogenic media (A) or in the presence of 100 ng/mL BMP-2 (B) and analyzed for enzymatic ALP activity represented as mean p-nitrophenyl phosphate (pNPP) content ± SEM. * represents p<0.05, ** represents <0.01, and *** represents p<0.001. **(a)** scaffold vs DBM; **(b)** scaffold vs GF; **(c)** DBM vs GF **(d)** scaffold OM vs BMP; **(e)** DBM OM vs BMP; **(f)** GF OM vs BMP; **(g)** DBM day 1 vs 3; **(h)** scaffold day 1 vs 3; **(i)** GF day 1 vs 3; **(j)** DBM day 3 vs 7; **(k)** scaffold day 3 vs 7; **(l)** GF day 3 vs 7 **(m)** DBM day 7 vs 15; **(n)** scaffold day 7 vs 15; **(o)** GF day 7 vs 15 **(p)** scaffold day 3 vs 15; **(q)** DBM day 3 vs 15; **(r)** GF day 3 vs 15

**Figure 6.**
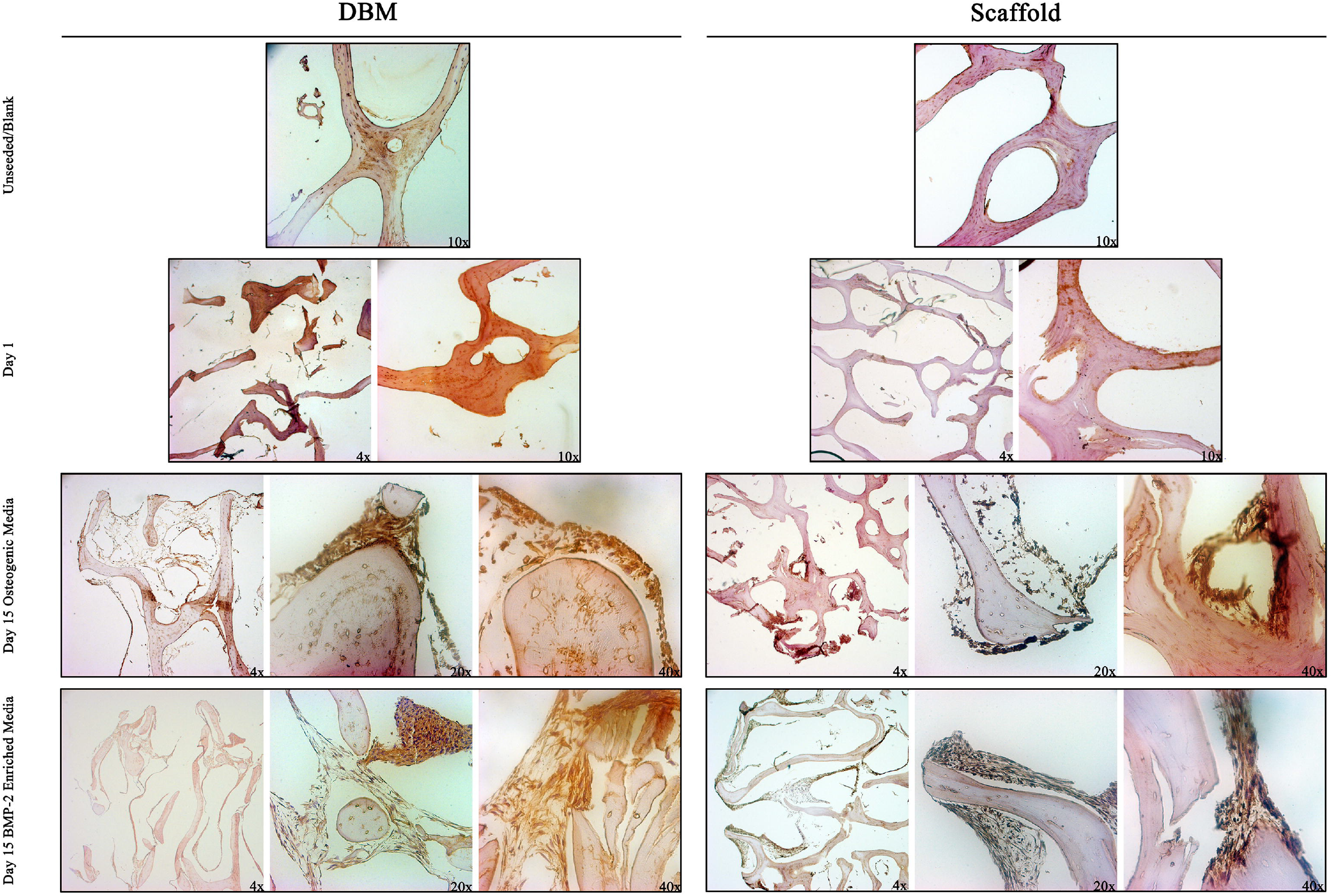
ALP expression increases over time in both DBM and scaffolds. DBM (left panels) or decellularized bone (right panels) scaffolds were sectioned unseeded, after 1-day culture of C2C12 cells or after 15 days in OM or OM with 100 ng/mL BMP-2. Sections were stained for ALP expression by immunohistochemistry and representative images are shown at 4X, 10X, 20X, or 40X.

To confirm the osteoinductive potential of the decellularized bone scaffolds, MC3T3-E1 preosteoblast cells were grown on the scaffolds. Compared to cells grown in a monolayer on collagen type I, MC3T3-E1 cells on the bone scaffolds expressed 2.5-fold higher *RANKL* and 2-fold higher *BMP-2* (Figure 7). Taken together, these results indicate the bone construct possesses osteoinductive potential *in vitro*.

**Figure 7.**
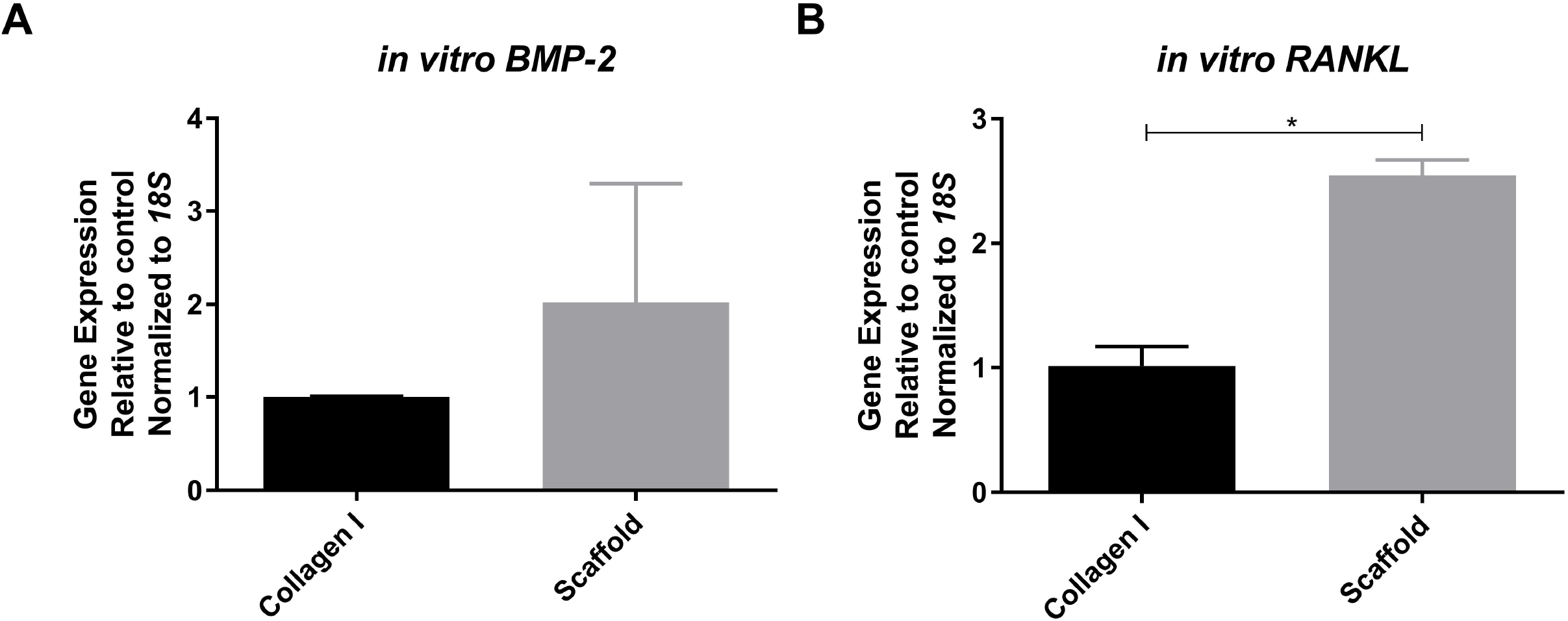
Scaffolds induce MC3T3-E1 osteogenic differentiation. MC3T3-E1 cells were seeded on 10 μg/mL collagen I (black bars) or decellularized bone scaffolds (gray bars) for one week. Gene expression of *BMP-2* (A) and *RANKL* (B) were measured and represented as mean fold change normalized to *18S* ± SEM.

### Implantation of Scaffolds in Syngeneic Mice Induces the Formation of a Bone Microenvironment

Based on the osteogenic and osteoinductive capabilities of the decellularized bone scaffold, we hypothesized that the seeded scaffold might be osteoconductive *in vivo*. MC3T3-E1 preosteoblasts were seeded on bone scaffolds and implanted subcutaneously in syngeneic mice. Unseeded scaffolds were implanted as controls. Upon removal, scaffolds were examined for changes in osteoblast differentiation gene expression. *In vivo* expression of *ALP* (3.3-fold), *BMP-2* (10.3-fold), and *BMP-7* (3.9-fold) increased within the pre-seeded scaffolds (Figure 8). *RANKL* gene expression was equal between groups (data not shown). To assess changes in the bone structure and measure possible bone formation, microCT analysis was performed on seeded (n=9) and unseeded (n=4) scaffolds (Figure 9). Increases in both bone volume fraction (BV/TV) and trabecular thickness (Tb.Th) were seen in the pre-seeded scaffolds; however, only Tb.Th reached significance (p=0.03). Paired t-tests showed significantly increased BV/TV (1.6-fold, p=0.0013) and Tb.Th (1.4-fold, p=0.0002) after explantation when both groups are combined (n=13) indicating new bone formation, regardless of cell seeding prior to implantation (Figure 9). After removal, scaffolds were section and analyzed for osteogenesis and angiogenesis. Pentachrome staining of the scaffolds displayed new bone formation within the pores of the scaffold in both groups (Figure 10A and 10C). To assess whether osteoclasts could be recruited to the scaffolds, staining for tartrate-resistant acid phosphatase (TRAP) was performed. Small TRAP-positive cells were seen within the pores and rare cells along the bone surface (Figure 10B and 10D). These cells likely represent osteoclast progenitor cells, although no mature osteoclasts were seen within the four-week time frame. Of note, pentachrome staining revealed apparent vascular-like structures (Figure 11A and 11C) within the scaffold. IHC analysis demonstrated positive CD31 staining of endothelial cells organized around a lumen and indicating angiogenesis within the decellularized bone scaffold (Figure 11B and 11D). These results demonstrated that the scaffold maintains osteoinductive potential following decellularization, due to its ability to recruit and stimulate cells down a bone forming lineage *in vivo*. Further, implantation of the scaffold in a syngeneic host, even without pre-osteoblast seeding, can generate a bone microenvironment.

**Figure 8.**
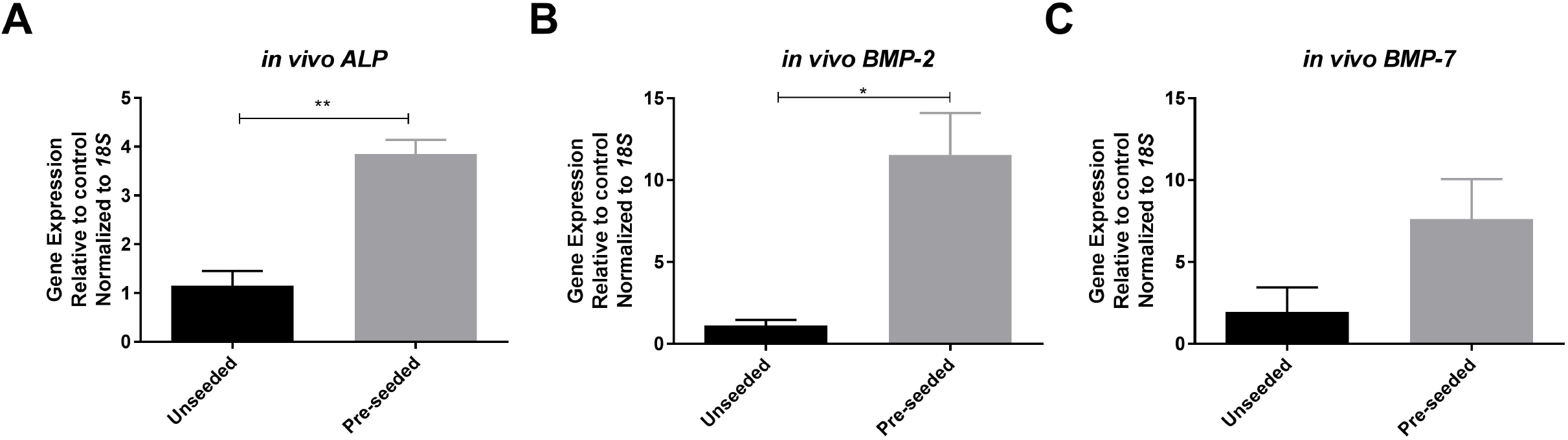
Implantation of scaffolds *in vivo* induces osteogenic gene expression. Bone scaffolds were seeded with MC3T3-E1 pre-osteoblast cells (gray bars) or left unseeded (black bars) and implanted subcutaneously in syngeneic mice. After 4 weeks scaffolds were removed, processed, and analyzed for gene expression of *ALP* (A), *BMP–2* (B), and *BMP-7* (C) represented as mean fold change normalized to *18S* ± SEM. * represents p<0.05.

**Figure 9.**
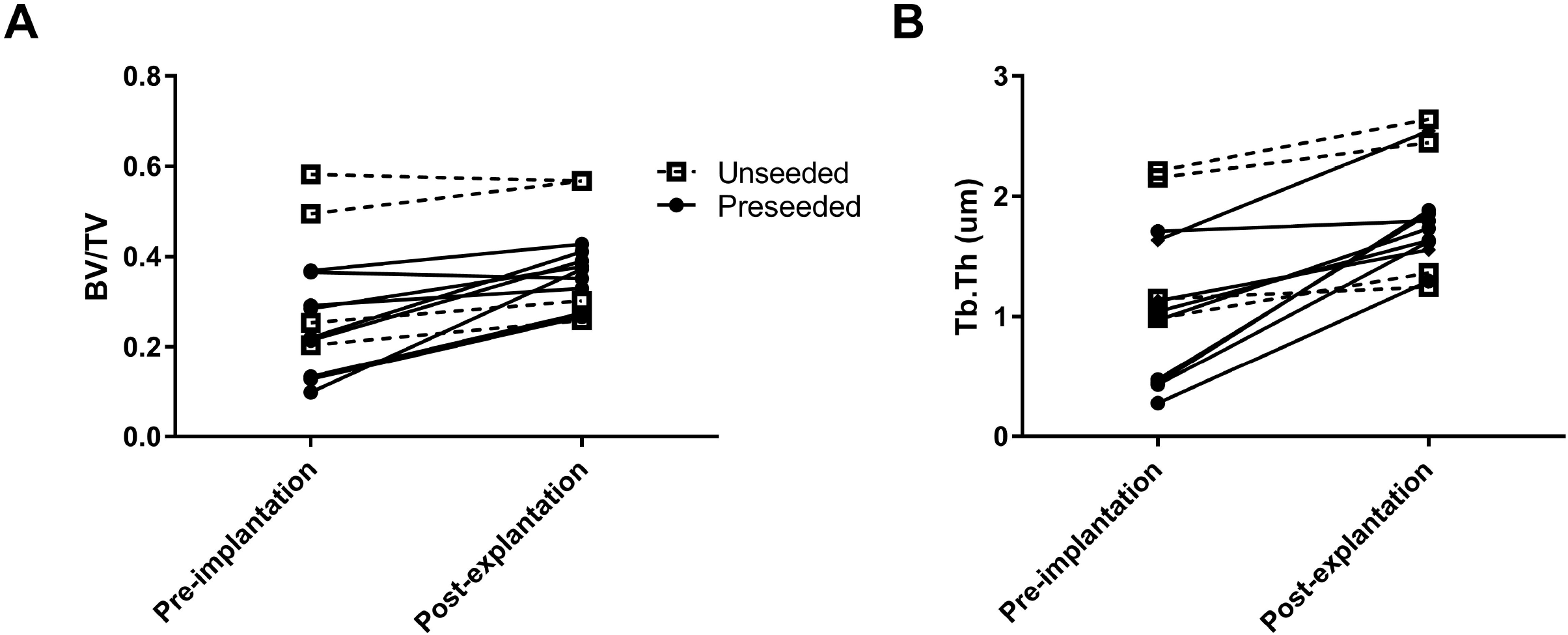
New bone formation occurs in implanted bone constructs. Bone scaffold structure was analyzed by microCT (pre-implantation) and then implanted subcutaneously in mice either unseeded (open squares) or pre-seeded with MC3T3-E1 pre-osteoblast cells (closed circles). Implants were removed after 4 weeks and scanned again by microCT (post-explantation). Bone volume ratio (BV/TV, A) and trabecular thickness (Tb.Th, B) were calculated and the change between individual scaffolds is shown.

**Figure 10.**
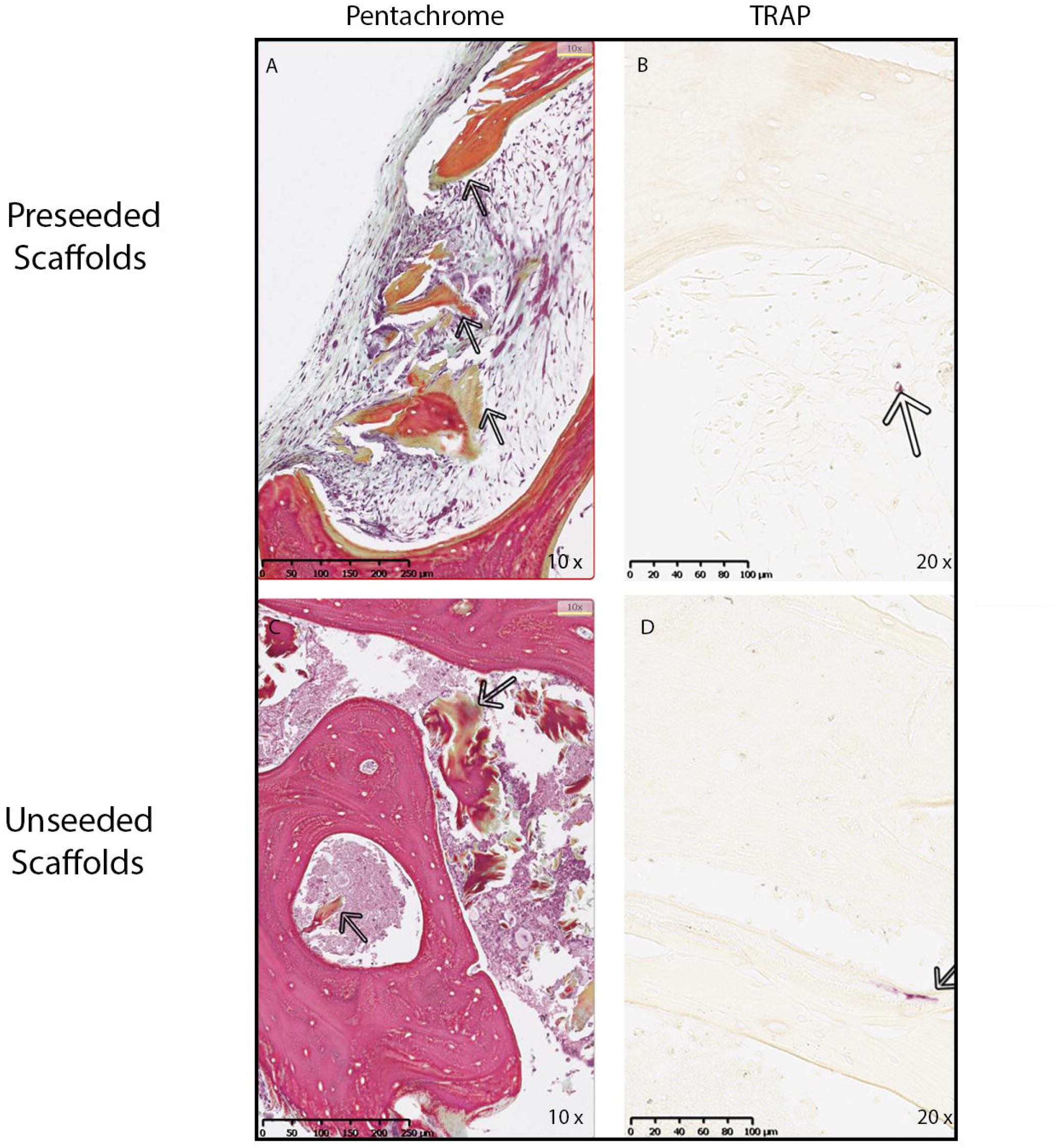
Implantation of bone scaffolds induces a functional bone microenvironment. MC3T3-E1 seeded and unseeded scaffolds were removed from mice after 4 weeks and processed for histology. Sections were stained with Movat’s pentachrome (A and C) to visualize the extracellular matrix deposition in the scaffolds. Arrows show the area of new bone formation in scaffold which stains green in representative photos captured at 10X. (B and D) Sections were also stained for osteoclasts using TRAP enzyme activity. Arrows point to purple staining demonstrating small TRAP-positive cells in the pores and along the bone surface in representative photos taken at 20X.

**Figure 11.**
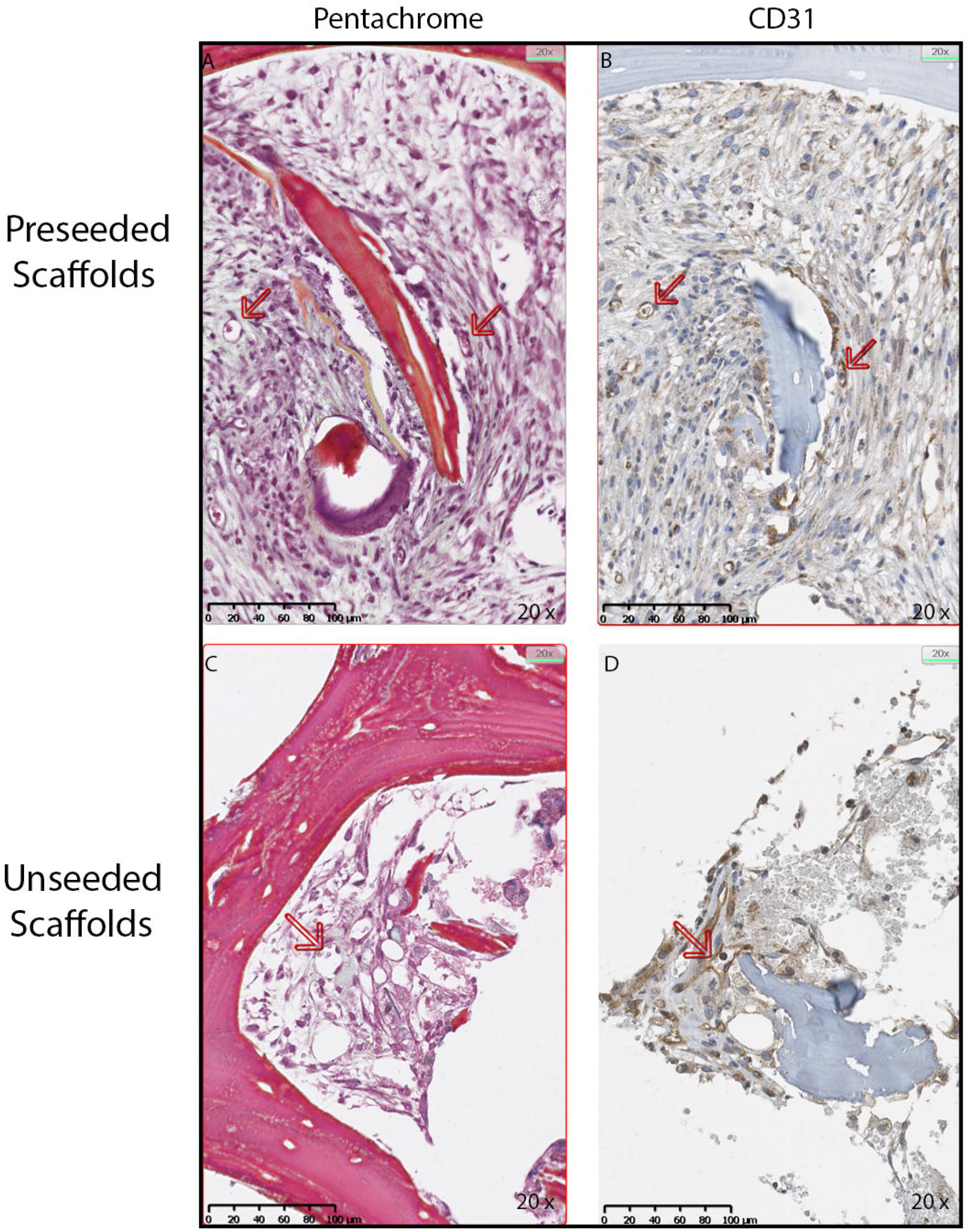
Angiogenesis occurs in the implanted bone scaffolds. MC3T3-E1 seeded and unseeded scaffolds were removed from mice after 4 weeks and processed for immunohistochemistry (IHC) for the endothelial marker CD31. In the representative micrographs at 20X, Movat’s pentachrome (A and C) demonstrate small vessels indicated with arrows, and corresponding IHC for CD31 confirms that these are endothelial cells (B and D).

## 5. Discussion/Conclusion

Previously, our laboratory established a decellularization and oxidation technique using PAA that removes 98% of DNA when applied to porcine bone [Bracey et al., 2018]. In this study, we demonstrate that this protocol preserves the native bone’s osteoinductive potential in a decellularized scaffold. The decellularized bone scaffold supported pre-osteoblast adhesion, survival, and proliferation comparable to DBM, a commercial product currently in clinical use with proven osteoinductive potential. Further, the bone scaffold supported the development of a bone microenvironment including osteogenesis, angiogenesis, and the recruitment of osteoclast precursors. Thus, the decellularized bone scaffold is osteoconductive and osteoinductive.

Control of osteogenesis *in vitro* and *in vivo* is integral to regenerative medicine applications in orthopaedic research. BMP-2 is one of the strongest stimulants of osteogenic differentiation in the pre-osteoblast cell lines used in our experiments [Katagiri et al., 1994; Han et al., 2003; Bormann et al., 2010; Qadir et al., 2015; Fu et al., 2017; Heo et al., 2018]. Concentrations as low as 100 ng/mL and 50 ng/mL were sufficient to promote osteogenic differentiation with increased ALP activity in MC3T3-E1 [Fu et al., 2017] and C2C12 [Han et al., 2003], respectively. However, few reports studied osteogenic differentiation of cells seeded onto xenograft-derived bone scaffolds [Arca et al., 2011; Hashimoto et al., 2011; Marcos-Campos et al., 2012; Kouroupis et al., 2013; Lu et al., 2013]. Hashimoto et al. demonstrated porcine hydroxyapatite contains osteoinductive properties and that these properties are maintained after processing [Hashimoto et al., 2011]. Similarly, Smith et al. found the osteoinductive properties were maintained in allografts following a decellularization and washing procedure [Smith et al., 2015]. However, Bormann et al. used a similar decellularization and oxidation protocol to ours with the addition of PAA on allografts and found the osteoinductive potential was not maintained [Bormann et al., 2010]. In the present study, we applied a decellularization and oxidation technique using PAA that removed 98% of the porcine DNA from the bone scaffolds [Bracey, 2017; Bracey et al., 2018]. Contrary to the findings by Borman et al. with human bone, our results demonstrate that the xenograft does indeed maintain osteoinductive potential after processing. C2C12 and MC3T3-E1 cells attached to the scaffold matrix, proliferated, and underwent osteogenic differentiation during the incubation period. The discrepancy between these studies outlines the variability between decellularization techniques as well as donor species. Bormann et al. reported that the donors ranged in both age (13-67 years) and gender, and ultimately concluded this could be a source of variability between the human-derived samples. These discrepancies may affect osteoinductive potential [Smith et al., 2017] and outline the importance of controlling environmental factors that may influence the quality of the donor bone, which is possible with the use of a xenograft.

Our *in vivo* microCT results demonstrated that there was equal osteogenesis identified in the pre-seeded and unseeded scaffolds. This finding suggests that the scaffold alone recruits the necessary cells for osteogenesis. Combining the microCT analysis and histologic assessment of the porcine bone scaffold *in vivo* demonstrated spontaneous new bone formation and angiogenesis. The identification of angiogenesis represents a critical finding due to the lack of vascularization being one of the major limitations associated with the use of tissue-engineered constructs during early bone regeneration [Muschler et al., 2004; Giannoni et al., 2010; Saran et al., 2014]. The presence of angiogenesis signifies graft-host integration by the induction of inflammatory cytokines as part of the normal healing process [Stegen et al., 2015]. It is reasonable to conclude that the presence of angiogenesis allowed for new bone formation due to the known importance angiogenesis has in bone repair and regeneration [Saran et al., 2014; Stegen et al., 2015]. Accordingly, Hirata et al. found that a BMP-2 soaked absorbable collagen sponge implanted in humans led to new bone formation lined by endothelial cells [Hirata et al., 2017]. Furthermore, Bhumiratana et al. implanted a clinically approved decellularized bovine trabecular bone seeded with adipose-derived stem cells into Yucatan minipig skull defects and concluded that angiogenesis and new bone formation occurred in parallel [Bhumiratana et al., 2016]. Thus, our decellularized bone scaffold represents a potential tissue-engineered bone microenvironment capable of both angiogenesis and osteogenesis. The angiogenesis demonstrated in these experiments may be due to the complex inflammatory response between the decellularized scaffold and M1 and M2 macrophages, and IL-4. This has been previously demonstrated within decellularized bone scaffold implants [Zheng et al., 2018], but is beyond the scope of this investigation.

There are limitations to our study. First, clinical translation of *in vitro* and animal experiments is limited. However, we believe these experiments are a necessary step to determine the properties of this bone scaffold after undergoing the decellularization and oxidation procedure. Second, our *in vivo* experiments involve an ectopic subcutaneous implantation model, rather than an orthotopic bone void filling model. However, the purpose of these experiments was solely to determine the osteoinductive potential of this scaffold in an *in vitro* and *in vivo* environment. Finally, a major limitation is using murine rather than human cell lines for these experiments, which limits immediate clinical translation. These cell lines, however, have been validated for the study of biomaterial osteoinductive potential previously [Qadir et al., 2015; Kanayama et al., 2017]. Despite these limitations, the decellularized porcine bone scaffold can be used in future experiments to assess its clinical and therapeutic utility.

Overall, our data demonstrate that a decellularization and oxidation technique applied to porcine metaphyseal bone preserves the osteoinductive potential of the bone. Previous literature identified that these properties are the most difficult to artificially create in tissue-engineered scaffolds and to maintain when processing bone scaffolds, therefore outlining the potential clinical impact of this construct. Future studies involving this xenograft will focus on placing the construct within a bone defect, identifying osseointegration, and comparing it to current standard treatments. These experiments will look at the effect of supplementing the scaffold with human mesenchymal stem cells, as a step towards clinical translation. Furthermore, an *in vivo* analysis of inflammatory markers to confirm that the bone scaffold has no increased reactivity when compared to currently used clinical implants for large bone defects should be performed. Further research will determine whether this decellularized bone scaffold capable of osteoinductivity and osteogenesis is fully osteoconductive and able to function as a substitute bone microenvironment.

## 8. Statements

### 8.1 Acknowledgments

We would like to thank Ms. Eileen Elsner, Ms. Jiaozhong Cia, and Dr. Lihong Shi for their help with tissue processing and animal handling throughout these experiments. We thank Brandi Bickford from the Virtual Microscopy Core for her assistance in slide scanning.

### 8.2 Statement of Ethics

All animal experiments conform to internationally accepted standards, and the study protocol was approved by the Wake Forest School of Medicine IACUC.

### 8.5 Author Contributions

Dr. Jinnah and Dr. Bracey had substantial contributions to the conception, acquisition, analysis, and interpretation of the work. Together they drafted the original manuscript and assisted with ongoing revisions. Dr. Willey contributed to analysis and interpretation of data and critical revisions to the manuscript. Dr. Seyler substantially contributed to the conception and design of the work and critical revisions to the manuscript. Dr. Hutchinson substantially contributed to the conception and design of the work and critical revisions to the manuscript. Dr. Whitlock substantially contributed to the conception and design of the work, interpretation of data, and critical revisions to the manuscript. Dr. Smith substantially contributed to the conception and design of the work, analysis of data, and critical revisions to the manuscript. Dr. Danelson substantially contributed to the analysis of the work and critical revisions to the manuscript. Dr. Emory substantially contributed to the interpretation of the work and critical revisions to the manuscript. Dr. Kerr substantially contributed to the conception and design of the work, analysis, interpretation of data, and critical revisions to the manuscript. All authors approved the final version of the manuscript for publication and agreed to be accountable for all aspects of the work.

## References

Albrektsson, T., C. Johansson (2001) Osteoinduction, osteoconduction and osseointegration. Eur Spine J 10 Suppl 2: S96–101.

Ansari, S., A. Moshaverinia, S.H. Pi, A. Han, A.I. Abdelhamid, H.H. Zadeh (2013) Functionalization of scaffolds with chimeric anti-BMP-2 monoclonal antibodies for osseous regeneration. Biomaterials 34(38): 10191–10198.

Araujo-Gomes, N., F. Romero-Gavilan, I. Garcia-Arnaez, C. Martinez-Ramos, A.M. Sanchez-Perez, M. Azkargorta, F. Elortza, J.J.M. de Llano, M. Gurruchaga, I. Goni, J. Suay (2018) Osseointegration mechanisms: a proteomic approach. Journal of biological inorganic chemistry: JBIC: a publication of the Society of Biological Inorganic Chemistry.

Arca, T., J. Proffitt, P. Genever (2011) Generating 3D tissue constructs with mesenchymal stem cells and a cancellous bone graft for orthopaedic applications. Biomed Mater 6(2): 025006.

Bhumiratana, S., J.C. Bernhard, D.M. Alfi, K. Yeager, R.E. Eton, J. Bova, F. Shah, J.M. Gimble, M.J. Lopez, S.B. Eisig, G. Vunjak-Novakovic (2016) Tissue-engineered autologous grafts for facial bone reconstruction. Science translational medicine 8(343): 343ra383.

Bormann, N., A. Pruss, G. Schmidmaier, B. Wildemann (2010) In vitro testing of the osteoinductive potential of different bony allograft preparations. Archives of orthopaedic and trauma surgery 130(1): 143–149.

Bracey, D.N. (2017) A Decellularized Porcine Xenograft-Derived Bone Scaffold for Clinical Use as a Bone Graft Substitute: A Critical Evaluation of Processing and Structure: Orthopaedic Surgery, Wake Forest University Graduate School of Arts & Sciences.

Bracey, D.N., T.M. Seyler, A.H. Jinnah, M.O. Lively, J.S. Willey, T.L. Smith, M.E. Van Dyke, P.W. Whitlock (2018) A Decellularized Porcine Xenograft-Derived Bone Scaffold for Clinical Use as a Bone Graft Substitute: A Critical Evaluation of Processing and Structure. Journal of functional biomaterials 9(3).

Calori, G.M., E. Mazza, M. Colombo, C. Ripamonti (2011) The use of bone-graft substitutes in large bone defects: any specific needs? Injury 42 Suppl 2: S56–63.

Campana, V., G. Milano, E. Pagano, M. Barba, C. Cicione, G. Salonna, W. Lattanzi, G. Logroscino (2014) Bone substitutes in orthopaedic surgery: from basic science to clinical practice. J Mater Sci Mater Med 25(10): 2445–2461.

Cooper, D.K.C., B. Ekser, A.J. Tector (2015) Immunobiological barriers to xenotransplantation. International journal of surgery 23(Pt B): 211–216.

De Long, W.G., Jr., T.A. Einhorn, K. Koval, M. McKee, W. Smith, R. Sanders, T. Watson (2007) Bone grafts and bone graft substitutes in orthopaedic trauma surgery. A critical analysis. The Journal of bone and joint surgery American volume 89(3): 649–658.

Fassbender, M., S. Minkwitz, M. Thiele, B. Wildemann (2014) Efficacy of two different demineralised bone matrix grafts to promote bone healing in a critical-size-defect: a radiological, histological and histomorphometric study in rat femurs. International orthopaedics 38(9): 1963–1969.

Feichtinger, G.A., T.J. Morton, A. Zimmermann, D. Dopler, A. Banerjee, H. Redl, M. van Griensven (2011) Enhanced reporter gene assay for the detection of osteogenic differentiation. Tissue engineering Part C, Methods 17(4): 401–410.

Fielding, G., S. Bose (2013) SiO2 and ZnO dopants in three-dimensionally printed tricalcium phosphate bone tissue engineering scaffolds enhance osteogenesis and angiogenesis in vivo. Acta biomaterialia 9(11): 9137–9148.

Franceschi, R.T., B.S. Iyer, Y. Cui (1994) Effects of ascorbic acid on collagen matrix formation and osteoblast differentiation in murine MC3T3-E1 cells. Journal of bone and mineral research: the official journal of the American Society for Bone and Mineral Research 9(6): 843–854.

Fu, C., X. Yang, S. Tan, L. Song (2017) Enhancing Cell Proliferation and Osteogenic Differentiation of MC3T3-E1 Pre-osteoblasts by BMP-2 Delivery in Graphene Oxide-Incorporated PLGA/HA Biodegradable Microcarriers. Scientific reports 7(1): 12549.

Giannoni, P., S. Scaglione, A. Daga, C. Ilengo, M. Cilli, R. Quarto (2010) Short-time survival and engraftment of bone marrow stromal cells in an ectopic model of bone regeneration. Tissue engineering Part A 16(2): 489–499.

Han, B., B. Tang, M.E. Nimni (2003) Quantitative and sensitive in vitro assay for osteoinductive activity of demineralized bone matrix. J Orthop Res 21(4): 648–654.

Hashimoto, Y., S. Funamoto, T. Kimura, K. Nam, T. Fujisato, A. Kishida (2011) The effect of decellularized bone/bone marrow produced by high-hydrostatic pressurization on the osteogenic differentiation of mesenchymal stem cells. Biomaterials 32(29): 7060–7067.

Heo, S.Y., S.C. Ko, S.Y. Nam, J. Oh, Y.M. Kim, J.I. Kim, N. Kim, M. Yi, W.K. Jung (2018) Fish bone peptide promotes osteogenic differentiation of MC3T3-E1 pre-osteoblasts through upregulation of MAPKs and Smad pathways activated BMP-2 receptor. Cell biochemistry and function 36(3): 137–146.

Hirata, A., T. Ueno, P.K. Moy (2017) Newly Formed Bone Induced by Recombinant Human Bone Morphogenetic Protein-2: A Histological Observation. Implant dentistry 26(2): 173–177.

Hsu, E.L., J.H. Ghodasra, A. Ashtekar, M.S. Nickoli, S.S. Lee, S.I. Stupp, W.K. Hsu (2013) A comparative evaluation of factors influencing osteoinductivity among scaffolds designed for bone regeneration. Tissue engineering Part A 19(15–16): 1764–1772.

Hupkes, M., A.M. Sotoca, J.M. Hendriks, E.J. van Zoelen, K.J. Dechering (2014) MicroRNA miR-378 promotes BMP2-induced osteogenic differentiation of mesenchymal progenitor cells. BMC Mol Biol 15(1): 1.

Iaquinta, M.R., E. Mazzoni, M. Manfrini, A. D’Agostino, L. Trevisiol, R. Nocini, L. Trombelli, G. Barbanti-Brodano, F. Martini, M. Tognon (2019) Innovative Biomaterials for Bone Regrowth. International journal of molecular sciences 20(3).

Kanayama, S., T. Kaito, K. Kitaguchi, H. Ishiguro, K. Hashimoto, R. Chijimatsu, S. Otsuru, S. Takenaka, T. Makino, Y. Sakai, A. Myoui, H. Yoshikawa (2017) ONO-1301 Enhances in vitro Osteoblast Differentiation and in vivo Bone Formation Induced by Bone Morphogenetic Protein. Spine.

Katagiri, T., A. Yamaguchi, M. Komaki, E. Abe, N. Takahashi, T. Ikeda, V. Rosen, J.M. Wozney, A. Fujisawa-Sehara, T. Suda (1994) Bone morphogenetic protein-2 converts the differentiation pathway of C2C12 myoblasts into the osteoblast lineage. J Cell Biol 127(6 Pt 1): 1755–1766.

Khan, S.N., F.P. Cammisa, Jr., H.S. Sandhu, A.D. Diwan, F.P. Girardi, J.M. Lane (2005) The biology of bone grafting. The Journal of the American Academy of Orthopaedic Surgeons 13(1): 77–86.

Kolambkar, Y.M., K.M. Dupont, J.D. Boerckel, N. Huebsch, D.J. Mooney, D.W. Hutmacher, R.E. Guldberg (2011) An alginate-based hybrid system for growth factor delivery in the functional repair of large bone defects. Biomaterials 32(1): 65–74.

Kouroupis, D., T.G. Baboolal, E. Jones, P.V. Giannoudis (2013) Native multipotential stromal cell colonization and graft expander potential of a bovine natural bone scaffold. Journal of orthopaedic research: official publication of the Orthopaedic Research Society.

Liu, G., J. Sun, Y. Li, H. Zhou, L. Cui, W. Liu, Y. Cao (2008) Evaluation of partially demineralized osteoporotic cancellous bone matrix combined with human bone marrow stromal cells for tissue engineering: an in vitro and in vivo study. Calcified tissue international 83(3): 176–185.

Livak, K.J., T.D. Schmittgen (2001) Analysis of relative gene expression data using real-time quantitative PCR and the 2(-Delta Delta C(T)) Method. Methods (San Diego, Calif) 25(4): 402–408.

Lu, X., J. Wang, B. Li, Z. Zhang, L. Zhao (2013) Gene expression profile study on osteoinductive effect of natural hydroxyapatite. J Biomed Mater Res A.

Marcos-Campos, I., D. Marolt, P. Petridis, S. Bhumiratana, D. Schmidt, G. Vunjak-Novakovic (2012) Bone scaffold architecture modulates the development of mineralized bone matrix by human embryonic stem cells. Biomaterials 33(33): 8329–8342.

Muschler, G.F., C. Nakamoto, L.G. Griffith (2004) Engineering principles of clinical cell-based tissue engineering. The Journal of bone and joint surgery American volume 86-A(7): 1541–1558.

Oryan, A., S. Alidadi, A. Moshiri, N. Maffulli (2014) Bone regenerative medicine: classic options, novel strategies, and future directions. Journal of orthopaedic surgery and research 9(1): 18.

Oryan, A., A. Kamali, A. Moshiri, M. Baghaban Eslaminejad (2017) Role of Mesenchymal Stem Cells in Bone Regenerative Medicine: What Is the Evidence? Cells, tissues, organs 204(2): 59–83.

Pierson, R.N., 3rd, A. Dorling, D. Ayares, M.A. Rees, J.D. Seebach, J.A. Fishman, B.J. Hering, D.K. Cooper (2009) Current status of xenotransplantation and prospects for clinical application. Xenotransplantation 16(5): 263–280.

Pina, S., R.F. Canadas, G. Jimenez, M. Peran, J.A. Marchal, R.L. Reis, J.M. Oliveira (2017) Biofunctional Ionic-Doped Calcium Phosphates: Silk Fibroin Composites for Bone Tissue Engineering Scaffolding. Cells, tissues, organs 204(3–4): 150–163.

Qadir, A.S., S. Um, H. Lee, K. Baek, B.M. Seo, G. Lee, G.S. Kim, K.M. Woo, H.M. Ryoo, J.H. Baek (2015) miR-124 negatively regulates osteogenic differentiation and in vivo bone formation of mesenchymal stem cells. J Cell Biochem 116(5): 730–742.

Roddy, E., M.R. DeBaun, A. Daoud-Gray, Y.P. Yang, M.J. Gardner (2018) Treatment of critical-sized bone defects: clinical and tissue engineering perspectives. European journal of orthopaedic surgery & traumatology: orthopedie traumatologie 28(3): 351–362.

Saran, U., S. Gemini Piperni, S. Chatterjee (2014) Role of angiogenesis in bone repair. Archives of biochemistry and biophysics 561: 109–117.

Seyler, T.M., D.N. Bracey, J.F. Plate, M.O. Lively, S. Mannava, T.L. Smith, J.M. Saul, G.G. Poehling, M.E. Van Dyke, P.W. Whitlock (2017) The Development of a Xenograft-Derived Scaffold for Tendon and Ligament Reconstruction Using a Decellularization and Oxidation Protocol. Arthroscopy: the journal of arthroscopic & related surgery: official publication of the Arthroscopy Association of North America and the International Arthroscopy Association 33(2): 374–386.

Shahi, M., M. Nadari, M. Sahmani, E. Seyedjafari, N. Ahmadbeigi, A. Peymani (2018) Osteoconduction of Unrestricted Somatic Stem Cells on an Electrospun Polylactic-Co-Glycolic Acid Scaffold Coated with Nanohydroxyapatite. Cells, tissues, organs 205(1): 9–19.

Shi, K., J. Lu, Y. Zhao, L. Wang, J. Li, B. Qi, H. Li, C. Ma (2013) MicroRNA-214 suppresses osteogenic differentiation of C2C12 myoblast cells by targeting Osterix. Bone 55(2): 487–494.

Shi, Q., Y. Li, J. Sun, H. Zhang, L. Chen, B. Chen, H. Yang, Z. Wang (2012) The osteogenesis of bacterial cellulose scaffold loaded with bone morphogenetic protein-2. Biomaterials 33(28): 6644–6649.

Shuang, Y., L. Yizhen, Y. Zhang, M. Fujioka-Kobayashi, A. Sculean, R.J. Miron (2016) In vitro characterization of an osteoinductive biphasic calcium phosphate in combination with recombinant BMP2. BMC oral health 17(1): 35.

Shui, W., W. Zhang, L. Yin, G. Nan, Z. Liao, H. Zhang, N. Wang, N. Wu, X. Chen, S. Wen, Y. He, F. Deng, J. Zhang, H.H. Luu, L.L. Shi, Z. Hu, R.C. Haydon, J. Mok, T.C. He (2013) Characterization of scaffold carriers for BMP9-transduced osteoblastic progenitor cells in bone regeneration. J Biomed Mater Res A.

Smith, C.A., T.N. Board, P. Rooney, M.J. Eagle, S.M. Richardson, J.A. Hoyland (2017) Human decellularized bone scaffolds from aged donors show improved osteoinductive capacity compared to young donor bone. PloS one 12(5): e0177416.

Smith, C.A., S.M. Richardson, M.J. Eagle, P. Rooney, T. Board, J.A. Hoyland (2015) The use of a novel bone allograft wash process to generate a biocompatible, mechanically stable and osteoinductive biological scaffold for use in bone tissue engineering. Journal of tissue engineering and regenerative medicine 9(5): 595–604.

Sondag, G.R., S. Salihoglu, S.L. Lababidi, D.C. Crowder, F.M. Moussa, S.M. Abdelmagid, F.F. Safadi (2013) Osteoactivin Induces Transdifferentiation of C2C12 Myoblasts into Osteoblasts. J Cell Physiol.

Stegen, S., N. van Gastel, G. Carmeliet (2015) Bringing new life to damaged bone: the importance of angiogenesis in bone repair and regeneration. Bone 70: 19–27.

Stiehler, M., F.P. Seib, J. Rauh, A. Goedecke, C. Werner, M. Bornhauser, K.P. Gunther, P. Bernstein (2010) Cancellous bone allograft seeded with human mesenchymal stromal cells: a potential good manufacturing practice-grade tool for the regeneration of bone defects. Cytotherapy 12(5): 658–668.

Thibault, R.A., L. Scott Baggett, A.G. Mikos, F.K. Kasper (2010) Osteogenic differentiation of mesenchymal stem cells on pregenerated extracellular matrix scaffolds in the absence of osteogenic cell culture supplements. Tissue engineering Part A 16(2): 431–440.

Torii, Y., K. Hitomi, N. Tsukagoshi (1994) L-ascorbic acid 2-phosphate promotes osteoblastic differentiation of MC3T3-E1 mediated by accumulation of type I collagen. Journal of nutritional science and vitaminology 40(3): 229–238.

Vadori, M., E. Cozzi (2015) The immunological barriers to xenotransplantation. Tissue antigens 86(4): 239–253.

Wancket, L.M. (2015) Animal Models for Evaluation of Bone Implants and Devices: Comparative Bone Structure and Common Model Uses. Vet Pathol 52(5): 842–850.

Wanschitz, F., E. Stein, W. Sutter, D. Kneidinger, K. Smolik, F. Watzinger, D. Turhani (2007) Expression patterns of Ets2 protein correlate with bone-specific proteins in cell-seeded three-dimensional bone constructs. Cells, tissues, organs 186(4): 213–220.

Whitlock, P.W., T.M. Seyler, G.D. Parks, D.A. Ornelles, T.L. Smith, M.E. Van Dyke, G.G. Poehling (2012) A novel process for optimizing musculoskeletal allograft tissue to improve safety, ultrastructural properties, and cell infiltration. The Journal of bone and joint surgery American volume 94(16): 1458–1467.

Whitlock, P.W., T.L. Smith, G.G. Poehling, J.S. Shilt, M. Van Dyke (2007) A naturally derived, cytocompatible, and architecturally optimized scaffold for tendon and ligament regeneration. Biomaterials 28(29): 4321–4329.

Yang, Q., J. Jian, S.B. Abramson, X. Huang (2011) Inhibitory effects of iron on bone morphogenetic protein 2-induced osteoblastogenesis. J Bone Miner Res 26(6): 1188–1196.

Yu, S., Q. Geng, J. Ma, F. Sun, Y. Yu, Q. Pan, A. Hong (2013) Heparin-binding EGF-like growth factor and miR-1192 exert opposite effect on Runx2-induced osteogenic differentiation. Cell Death Dis 4: e868.

Zimmermann, G., A. Moghaddam (2011) Allograft bone matrix versus synthetic bone graft substitutes. Injury 42 Suppl 2: S16–21.

